# Interaction with IP6K1 supports pyrophosphorylation of substrate proteins by the inositol pyrophosphate 5-IP7

**DOI:** 10.1101/2023.12.10.570918

**Authors:** Aisha Hamid, Jayashree S. Ladke, Akruti Shah, Monisita Pal, Shubhra Ganguli, Arpita Singh, Rashna Bhandari

## Abstract

Inositol pyrophosphates (PP-IPs) are a sub-family of water soluble inositol phosphates that possess one or more diphosphate groups. PP-IPs can transfer their β-phosphate group to a phosphorylated Ser residue to generate pyrophosphorylated Ser. This unique post-translational modification occurs on Ser residues that lie in acidic stretches within an intrinsically disordered protein sequence. Serine pyrophosphorylation is dependent on the presence of Mg^2+^ ions, but does not require an enzyme for catalysis. The mechanisms by which cells can regulate this enzyme-independent modification are still unknown. Here, we show that IP6K1, an enzyme responsible for the synthesis of the PP-IP 5-IP7, interacts with several proteins that undergo 5-IP7 mediated pyrophosphorylation, and with CK2, a protein kinase that phosphorylates Ser residues prior to pyrophosphorylation. We characterized the interaction of IP6K1 with AP3B1, the β subunit of the AP3 adaptor protein complex, which is a known pyrophosphorylation substrate. We observe the formation of a protein complex between IP6K1, AP3B1, and the catalytic α-subunit of CK2, and show that disrupting IP6K1 binding to AP3B1 lowers its in vivo pyrophosphorylation. We propose that assembly of a substrate-CK2-IP6K complex would allow for coordinated pre-phosphorylation and pyrophosphorylation of the target serine residue, and provide a mechanism to regulate this enzyme-independent modification.

## 1. Introduction

Inositol pyrophosphates (PP-IPs) are a sub-family of water soluble inositol phosphates that contain one or more energy-rich diphosphate moieties. Found in all eukaryotic organisms, the best studied PP-IPs in mammals are 5-IP7 (5-diphosphoinositol pentakisphosphate or 5PP-IP5) and 1,5-IP8 (bisdiphosphoinositol tetrakisphosphate or 1,5(PP)2-IP4). The synthesis of 5-IP7 from IP6 (inositol hexakisphosphate) is catalyzed by IP6 kinases, of which there are three isoforms in mammals – IP6K1, IP6K2 and IP6K3 [1-3]. 5-IP7 is a substrate for two mammalian PP-IP5 kinases, which catalyze the synthesis of 1,5-IP8 [4]. Designated as metabolic messengers, PP-IPs are known to influence several cellular functions including phosphate homeostasis, apoptosis, vesicle trafficking, insulin secretion, ribosome biogenesis, and DNA repair [5-8].

PP-IPs selectively regulate protein function using two molecular mechanisms - allosteric non-covalent binding, and covalent protein pyrophosphorylation [6, 9]. PP-IP mediated serine pyrophosphorylation is a unique enzyme-independent post translational modification. During pyrophosphorylation of a substrate protein, PP-IP transfers its β-phosphate to a pre-phosphorylated serine residue located within an intrinsically disordered acidic serine sequence motif, in the presence of magnesium cations [9-11]. Although pre-phosphorylation of the target serine residues is brought about by acidophilic Ser/Thr kinases, phosphotransfer of the β-phosphate from the donor PP-IP to the acceptor phosphoserine residue can take place without any enzyme. Using radiolabeling methods, 5-IP7 mediated pyrophosphorylation has been demonstrated individually for *S. cerevisiae* proteins Nsr1, Srp40, Rpa190, Rpa43, Rpa34, and Gcr1, and mammalian proteins AP3B1, DIC2C, MYC, NOLC1, TCOF and UBF1 [10-17]. Recently, 148 sites of pyrophosphorylation were identified in 71 human proteins by a mass-spectrometry based proteomics analysis, of which the nucleolar proteins NOLC1 and TCOF possess the most number of pyrophosphosites [17]. It was shown that endogenous pyrophosphorylation of these proteins is indeed dependent on 5-IP7, as co-expression of active IP6K1 led to increased pyrophosphorylation of these proteins, whereas inactive IP6K1 did not have any effect.

The biological consequences of pyrophosphorylation by PP-IPs have been reported in the case of a few protein substrates. Pyrophosphorylation of the budding yeast transcription factor Gcr1 destabilizes its interaction with its partner Gcr2, stalling the transcription of essential glycolytic enzymes and lowering the rate of glycolysis [13]. Also in budding yeast, 5-IP7 has been shown to pyrophosphorylate three subunits in RNA polymerase I – Rpa190, Rpa43, and Rpa34. Yeast lacking Kcs1, the enzyme responsible for 5-IP7 synthesis, exhibited a reduced rate of rRNA synthesis, leading to defects in ribosome biogenesis and a reduction in protein translation [16]. The inhibition of IP6 kinases also resulted in reduced rRNA transcription in mammalian cells [17]. This phenotype may be attributed to the high level of endogenous pyrophosphorylation of nucleolar proteins in the fibrillar center of the nucleolus where rDNA transcription takes place. Pyrophosphorylation of three different mammalian proteins has been shown to influence their function by regulating their interaction with other proteins. 5-IP7 mediated pyrophosphorylation at Serine51 on the dynein-intermediate chain (DIC2C) promotes its interaction with the p150Glued subunit of dynactin, enhancing recruitment of the dynein motor to vesicles [15]. The oncoprotein MYC undergoes pyrophosphorylation in its central PEST domain [14]. This modification is essential for binding of the MYC PEST domain to the E3 ubiquitin ligase FBW7, which in turn promotes MYC polyubiquitination and turnover. Pyrophosphorylation of AP3B1, the β subunit of the AP3 adaptor protein complex disrupts its binding to the microtubule plus-end directed kinesin motor protein KIF3A [12]. The AP3 complex and KIF3A promote the intracellular trafficking and subsequent release of HIV1 Gag protein. 5-IP7 mediated pyrophosphorylation of AP3B1 therefore lowered the release of HIV-1 virus-like particles from mammalian cells.

Although the biological functions of protein pyrophosphorylation by PP-IPs are now well established, our understanding of the molecular details of this phosphotransfer is still nascent. Enzyme-independent phosphate transfer from PP-IP to the target protein is supported by a high concentration of Mg^2+^ in the buffer, and increasing temperature [9-11]. Other essential factors are the acidic amino acid residues flanking the target phosphoserine residue, and the presence of this target sequence within an intrinsically disordered region of the substrate protein [9]. It is speculated that disorder in the substrate protein facilitates its interaction with the Mg^2+^-PP-IP complex, but in the absence of detailed biochemical and structural characterization, the mechanism of the phosphate transfer from PP-IP to phosphoserine is not clear.

IP6 and IP7 have been shown to bind proteins with comparable specificity and affinity [18, 19], as these molecules are similar in their structure and charge, barring the presence of a β phosphate in IP7. The presence of excess IP6 can inhibit in vitro phosphotransfer from 5-IP7 to a protein substrate [10], suggesting that pyrophosphorylation substrate proteins can bind IP6 in addition to PP-IPs. As cells are reported to harbour a 5- to 50-fold higher intracellular concentration of IP6 compared with PP-IPs [20], binding of IP6 to a target protein may inhibit its intracellular pyrophosphorylation by PP-IPs. The data presented in our study addresses this concern. Our analysis of the protein interactome of IP6K1 revealed that it interacts with several pyrophosphorylation substrate proteins, and with the protein kinase CK2, which is primarily responsible for pre-phosphorylation of serine residues that are targeted for pyrophosphorylation by 5-IP7 [11]. We show that IP6K1 forms a multi-protein complex with the pyrophosphorylation substrate protein AP3B1 and the pre-phosphorylating kinase CK2. Binding of IP6K1 with AP3B1 is primarily via an intrinsically disordered region in the N-terminal lobe of IP6K1. Intracellular over-expression of this IP6K1 fragment disrupts the interaction of IP6K1 with AP3B1, and lowers endogenous AP3B1 pyrophosphorylation. Binding of the 5-IP7 synthesizing IP6 kinase with the pyrophosphorylation substrate protein may provide a mechanism to increase the local concentration of 5-IP7 and counter any inhibition of pyrophosphorylation by IP6. Formation of a substrate-CK2-IP6K complex would also facilitate coordinated pre-phosphorylation and pyrophosphorylation of the target serine residue, providing cells with a mechanism to control this enzyme-independent modification.

## 2. Materials and Methods

### 2.1 Reagents

The primary antibodies used in this study for immunoblotting (IB) or immunoprecipitation (IP), along with antibody dilution for each application, and supplier (including catalogue number), were as follows: anti-IP6K1 (Sigma-Aldrich, HPA040825; IB, 1:3000; Genetex, GTX103949; IB 1:3,000); anti-GAPDH (Sigma-Aldrich, G8795; IB, 1:10,000); anti-α-TUBULIN (Sigma-Aldrich, T9026; IB, 1:10,000); anti-V5 tag (Thermo Fisher Scientific, R960-25; IB, 1:7000; IP, 1 μg,); anti-HA (Sigma-Aldrich, H6908; IP, 1 μg, IB: 1:7000); anti-FLAG (Sigma-Aldrich, F1804; IB, 1:10,000; IP, 1 μg); anti-myc tag (Sigma-Aldrich, M4439; IB, 1:10,000; IP, 1 μg;), anti-AP3B1 (Bethyl labs 50-157-0195, IB, 1:2000); anti-TCOF (Abclonal A2512 IB, 1:1000); anti-UBF1 (Santa Cruz, sc-13125 IB, 1:500). For immunoprecipitation of endogenous IP6K1, an in-house antibody generated against the N-terminal region of IP6K1 was used [21]. PVDF membrane for protein transfer, Glutathione-sepharose beads, Protein A-sepharose beads, Protein G-sepharose beads, Streptavidin-sepharose beads, and ECL prime chemiluminescence substrate were procured from GE Healthcare. The Gateway Clonase enzymes were purchased from Thermo Fisher Scientific. NuPAGE 4-12% Bis-Tris gels, 20X MES running buffer, and 4X LDS sample buffer were purchased from Thermo Fisher Scientific. [γ-^32^P]ATP (LCP-101) was procured from JONAKI/BRIT. *myo*-2-[^3^H] inositol (15-20 Ci/mmol) (ART 0116B) was procured from American Radiolabeled Chemicals. Ultima-Flo AP (6013599) was purchased from Perkin-Elmer. Casein Kinase II (CK2 holoenzyme) (P6010) was purchased from New England Biolabs.

### 2.2 Plasmids

Human IP6K1 cDNA (GenBank ID NM_001242829.2) and catalytically inactive IP6K1 (K226A) mutant were subcloned in pCDNA3.1 as previously described [21]. N-terminally myc-tagged human NOLC1 cDNA (GenBank ID NM_001284388.2) expression plasmid was procured from Sino Biologicals (HG16317-NM). cDNA encoding human IP6K1 domains, i.e., FL, N-lobe, C-lobe, IDR-1, IDR-2, PDKG motif with IP helices, and IDR-1 with C-lobe, were cloned into the N-terminal SFB-tag destination vector, using the Gateway cloning strategy (Thermo Fisher Scientific). Mouse full length MYC cDNA with a C-terminal V5 tag was cloned in pCDNA 3.1 as previously described [14]. pZW12 CK2β was a gift from David Litchfield (Addgene plasmid 27088; GenBank ID NM_001320.7). Human AP3B1 (GenBank ID NM_003664.5) cDNA (a gift from Solomon Snyder, Johns Hopkins School of Medicine, Baltimore, USA) was subcloned into SalI and NotI restriction enzyme sites into the pCMV_Myc plasmid vector to express myc epitope-tagged AP3B1. Human CK2α cDNA in the pCMV-HA vector was a gift from Solomon Snyder. Plasmid expressing HA-tagged CK2α (K68M) was subcloned from pGV15-CK2α-K68M (a gift from David Litchfield, Addgene plasmid 27089) into SalI and NotI restriction enzyme sites in the pCMV-HA vector.

### 2.3 Cell culture and transfection

The HEK293T cell line was obtained from the laboratory of Solomon Snyder at Johns Hopkins School of Medicine, Baltimore, USA, and was authenticated by Lifecode Technologies Private Limited, New Delhi, India. Short tandem repeat profiling for authentication showed a 94.1% match with the ATCC reference genotype (https://www.atcc.org/products/crl-3216). HEK293T cells were grown in a humidified incubator with 5% CO_2_ at 37°C in Dulbecco’s modified Eagle’s medium (DMEM) supplemented with 10% fetal bovine serum, 1 mM L-glutamine, 100 U/mL penicillin, and 100 μg/mL streptomycin. For transfection of HEK293T cells, polyethylenimine (PEI) (Polysciences, 23966) was used at a ratio of 1:3 (DNA:PEI). All plasmids used for transfection were purified using the Plasmid Midi kit (Qiagen). Cells were harvested 36-48 h post-transfection for further analyses.

### 2.4 Tandem affinity purification and mass spectrometry analysis

HEK293T cells transiently transfected to express either SFB-tagged GFP or SFB-tagged IP6K1 were lysed in NETN buffer (20 mM Tris-HCl, pH 8.0, 100 mM NaCl, 1 mM EDTA, 0.5% Nonidet P-40) containing protease inhibitor cocktail (Sigma-Aldrich, P8340) and phosphatase inhibitor cocktail (Sigma-Aldrich, P5762), at 4°C for 60 min. Cell debris were removed by centrifugation at 14000 x *g* for 10 min, and cell lysates were incubated with streptavidin-sepharose beads (Cytiva Life Sciences) for 1 h at 4°C with end-over-end rotation. The bound protein complexes were washed thrice with NETN buffer and then eluted with 2 mg/mL biotin (Merck) for 90 min at 4°C. The eluates were incubated with S-protein agarose beads (Novagen) for 1 h at 4°C and then washed thrice with NETN buffer. Proteins bound to S-protein agarose beads were boiled in 2x SDS sample buffer for 5 min, loaded on a 12% SDS polyacrylamide gel, allowed to run into the resolving gel up to 1 cm, and visualized by Coomassie Brilliant Blue staining. All proteins in the sample were excised in one gel slice and sent for mass spectrometry analysis to Taplin Biological Mass Spectrometry Facility at Harvard University, USA. Briefly, gel pieces were subjected to in-gel digestion with sequencing-grade trypsin (Promega), and the extracted peptides were resolved by reverse phase HPLC. Eluted peptides were subjected to electrospray ionization (ESI) and then allowed to enter an LTQ Orbitrap Velos Pro ion-trap mass spectrometer (Thermo Fisher Scientific). Peptides were detected, isolated, and fragmented to produce a tandem mass spectrum of specific fragment ions for each peptide. Peptide sequences (and hence protein identity) were determined by matching protein databases from UniProt (https://www.uniprot.org/taxonomy/10090) with the acquired fragmentation pattern by the software program, Sequest (Thermo Fisher Scientific). Mass spectromery data were submitted to the MassIVE repository, a full member of the Proteome Xchange Consortium. Data can be accessed via the URL <https://massive.ucsd.edu>, with the data set identifier MassIVE MSV000092218. The total peptide number for each protein, which reflects relative protein abundance in the sample, was compared between the test (SFB-IP6K1) and control (SFB-GFP) samples using the CRAPome tool (https://reprint-apms.org)[22]. Empirical Fold Change Score (FCA and FCB) were used to compare the enrichment of proteins in the bait (SFB-IP6K1) over user control (SFB-GFP) in replicates. A fold change score (FC-B) of 1.4 was used as a cut-off score to select genuine interactors of IP6K1. The selected proteins were examined for Gene Ontology (GO) term enrichment using the Functional Annotation Clustering tool at Database for Annotation, Visualization and Integrated Discovery (DAVID) v6.7 (https://david.ncifcrf.gov) [23].

### 2.5 Co-immunoprecipitation assay

HEK293T cells were collected 48 h post transfection, lysed for 1 h at 4°C in lysis buffer (50 mM HEPES, pH 7.4, 100 mM NaCl, 1 mM EDTA, 0.5 % Nonidet P-40, protease inhibitor cocktail (Sigma-Aldrich, P8340) and phosphatase inhibitor cocktail (Sigma-Aldrich, P5762). The lysates were incubated with specific antibody overnight at 4°C with end-over-end mixing. The complex was pulled down using Protein A or Protein G-sepharose beads (GE Healthcare) pre-equilibrated in lysis buffer for 2 h. The beads were washed thrice with lysis buffer followed by boiling in 1 x Laemmli buffer. The samples were analysed by immunoblotting with primary antibodies (details are provided above). For SFB-tagged protein pull-down, streptavidin-sepharose beads were added to the cell lysate for 2 h at 4°C followed by washing and boiling in 1 x Laemmli buffer, and analysis by immunoblotting. UVITEC Alliance Q9 documentation system or the GE ImageQuant LAS 500 imager were used for chemiluminescence detection. Fiji ImageJ software was used for densitometry based quantitative analysis. For quantification of protein levels, band intensities were normalized to that of the loading control. For quantification of the extent of co-immunoprecipitation, band intensities of the co-precipitating proteins were normalized to intensity of the immunoprecipitated protein.

### 2.6 Purification and pull-down with GST fusion proteins

GST-tagged IP6K1 and CK2α were expressed in *E. coli* BL21 (DE3) strain and affinity chromatography was performed using standard protocols. Cells were grown in LB medium, and protein expression was induced with the addition of 0.5 mM ITPG overnight at 18°C. Cultures were pelleted by centrifuging at 6000 x *g* and pellets were lysed by sonication in ice-cold buffer A (20 mM HEPES pH 6.8, 100 mM NaCl, 2 mM EDTA, and 5 mM DTT). The lysates were centrifuged at 18000 x *g* and the supernatant was incubated with equilibrated glutathione-sepharose beads at 4°C for 2 h. Beads were washed twice with Buffer C (20 mM HEPES pH 6.8, 500 mM NaCl, 2 mM EDTA, 1% Triton X-100) and twice with Buffer B (20 mM HEPES pH 6.8, 100 mM NaCl, 2 mM EDTA, 1% Triton X-100). The beads were resuspended in an equal volume of 1X PBS and 20 μL slurry was analysed by SDS-PAGE to determine the amount of purified protein, with a known amount of BSA for comparison. Purified GST, or GST-tagged IP6K1 or CK2α immobilized on glutathione-sepharose beads were incubated with HEK293T cell lysates in buffer containing 50 mM HEPES-KOH pH 7.4, 100 mM NaCl, 5 mM MgCl_2_, 0.3% Triton X-100, and protease and phosphatase inhibitor cocktails, for 2 h at 4°C. The interacting proteins were identified by immunoblotting.

### 2.7 Analysis of cellular inositol pyrophosphates

IP6K1 knockout HEK293T cells, transfected to express either active or catalytically inactive IP6K1, were labeled with [^3^H]-inositol as described earlier [24, 25]. Cells seeded in 60 mm dishes in normal growth medium were allowed to attain 30% confluence and then transferred to inositol-free DMEM (MP Biomedicals, D9802-06.25) containing 10% dialysed fetal bovine serum, and labelled with 30 μCi myo-2-[^3^H] inositol for 2.5 days. The media was removed and fresh media containing myo-2-[^3^H] inositol (30 μCi) was added for another 2.5 days. At the end of 5 days, when isotopic labeling is achieved, cells were washed and collected by scraping in chilled PBS. From the cell pellet, soluble inositol phosphates were extracted by the addition of 350 μL extraction buffer (0.6 M HClO4, 2 mM EDTA, 0.2 mg/mL phytic acid (Sigma-Aldrich, P8810) on ice for 15-20 min, followed by centrifugation at 21,000 x *g* for 10 min. The supernatant containing soluble inositol phosphates was collected, and lipid inositols in the pellet were extracted with 1 mL lipid extraction buffer (0.1 N NaOH and 0.1% Triton X-100) at room temperature with end-over-end mixing for 4-5 h. The soluble inositol phosphate extract was mixed with ∼120 μL neutralisation solution (1 M K_2_CO_3_ and 5 mM EDTA). Tubes were left open on ice for 1 h, followed by centrifugation at 21,000 x *g* for 10 min at 4°C. The extracted inositol phosphates were resolved by HPLC (5125 HPLC pumps, Waters) on a Partisphere SAX column (4.6 mm × 125 mm, HiChrome) using a gradient of Buffer A (1 mM EDTA) and Buffer B (1mM EDTA and 1.3 M (NH4)2HPO4 (pH 3.8)] as follows - 0-5 min, 0% B; 5-10 min, 0-20% B; 10-70 min, 20-100% B; 70-80 min, 100% B. 1 mL fractions containing soluble inositol phosphates were mixed with 3 mL scintillation cocktail (Ultima-Flo AP) and counted for 5 min in a liquid scintillation counter (Tri-Carb 2910 TR, Perkin Elmer). The lipid inositol extract was counted in a liquid scintillation counter, and the soluble inositol phosphate count in each fraction was normalized to this.

### 2.8 Protein phosphorylation and pyrophosphorylation

For the phosphorylation reactions, myc-tagged AP3B1 immunoprecipitated and immobilised on Protein G-sepharose beads was incubated with CK2 (250 units; New England Biolabs) for 30 min, in the presence of protein kinase buffer (New England Biolabs) with 3 μCi [γ-^32^P]ATP, and 500 μM Mg^2+^-ATP. Beads were washed with ice-cold PBS and eluted in 1X LDS sample buffer followed by boiling at 95°C for 5 min. Proteins were resolved by a 4-12% NuPAGE Bis-Tris gel and transferred to a PVDF membrane. Blots were dried and subjected to autoradiography, followed by immunoblotting to detect myc-tagged AP3B1.

The synthesis of 5[β-^32^P]IP7, and pyrophosphorylation of proteins expressed in HEK293T cells using radiolabeled 5-IP7 have been described previously [14, 15, 17, 26]. Briefly, myc-tagged AP3B1 expressed transiently in HEK293T cells was immunoprecipitated using a myc-tag antibody and immobilised on protein G-sepharose beads. The immunoprecipitated complexes were resuspended in pyrophosphorylation buffer (25 mM HEPES-KOH pH 7.4, 50 mM NaCl, 6 mM MgCl_2_, 1 mM DTT) in the presence of 3 μCi 5[β-^32^P]IP7, and incubated at 37°C for 15 min. The beads were boiled in 1X LDS sample buffer at 95°C for 5 min, resolved by 4-12% NuPAGE Bis-Tris gel and transferred to a PVDF membrane. The blots were exposed for about ∼5 days to a phosphorimager screen. Phosphorylation and pyrophosphorylation of AP3B1 were detected using a phosphorimager (Typhoon FLA-9500). The ratio of radiolabelled protein to the total immunoprecipitated protein was quantified by densitometry analysis using the Fiji ImageJ software.

## 3. Results

### 3.1. Identification of the IP6K1 protein interactome

To identify the specific interacting partners of IP6K1, SFB-tagged IP6K1 was over-expressed in HEK293T cells and subjected to affinity purification under non-denaturing conditions, followed by mass spectrometry. The protein interactome of SFB-tagged GFP served as a negative control (Table S1). Pull-down and mass spectrometry analysis for both IP6K1 and GFP was conducted in replicate sets (Figure 1A). A total of 414 proteins were found to uniquely bind IP6K1, of which 57 were present in both replicate sets (Figure 1A). This data was subjected to the CRAPome analysis pipeline (reprint-apms.org/) to distinguish between authentic and contaminant interactors by comparing the total peptide number for each interacting protein bound to the test sample (IP6K1) with the control sample (GFP) (Table S2). In the list of specific IP6K1 interacting proteins, we observed DDB1 and EIF4G1, which are experimentally verified binding partners of IP6K1, contributing to its role in CRL4 signalosome assembly and P-body formation respectively [21, 27]. Additional proteins in the IP6K1 specific interactome have previously been identified in high-throughput studies on protein-protein interactions [28, 29]. We set the stringent fold change FC-B score [22] to 1.4 to accommodate the verified IP6K1 interactor EIF4G1 [21], and subjected 179 IP6K1 interacting proteins that fall in this range to Gene Ontology (GO) term enrichment analysis using the DAVID tool (https://david.ncifcrf.gov) [23]. IP6K1 interacting partners were found to be involved in several biological processes including translation, mRNA splicing, and DNA repair (Figure 1B).

**Figure 1.**
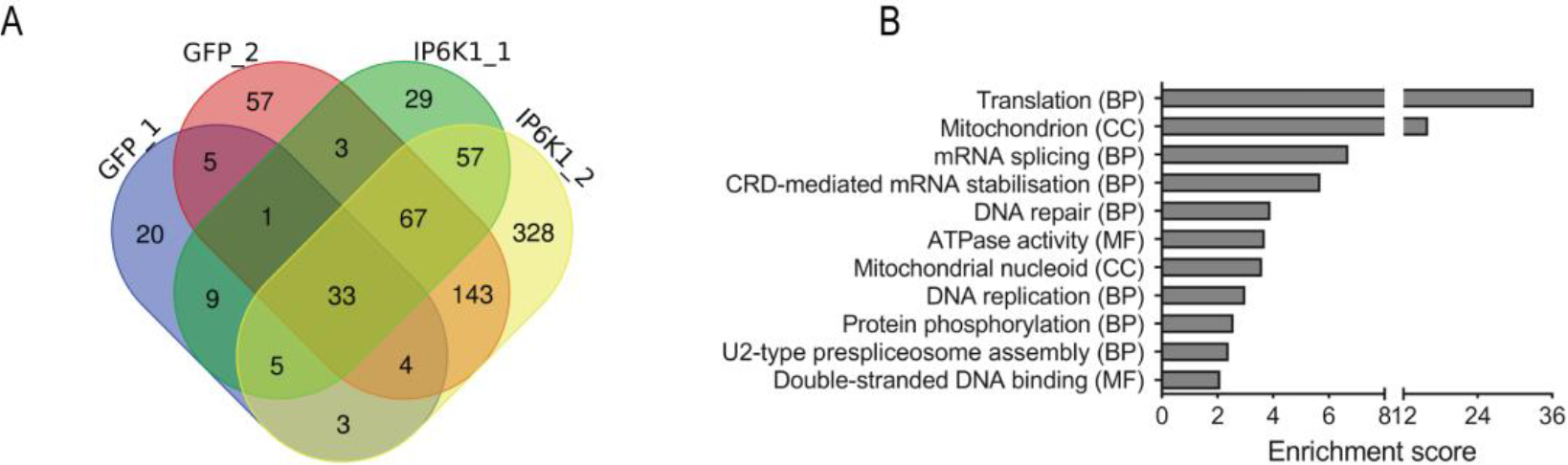
Analysis of the IP6K1 protein interactome. **(A)** Venn diagram depicting the number of proteins identified in each replicate of SFB-tagged GFP (control) or SFB-tagged IP6K1. A total of 414 proteins that interacted specifically with IP6K1 were absent from the GFP pull-down. **(B)** The total peptide count for each protein bound to replicate test (IP6K1) and control (GFP) samples was analyzed using the CRAPome tool; proteins that show a fold change (FC-B) score of ≥ 1.4 were subjected to Functional Annotation Clustering of Gene Ontology (GO) terms using the DAVID tool (Table S3). The graph represents the clusters of Cellular Component (CC), Biological Process (BP) and Molecular Function (MF) GO terms with a group enrichment score ≥ 2 (P ≤ 0.01).

Interestingly, the IP6K1 interactome contained 5-IP7 pyrophosphorylation substrates including AP3B1, NOLC1, TCOF and UBF1 (Table 1 and Table S2) [10-12, 17]. Among the interactors we also observed the catalytic α and α’ (CSK21 and CSK22), and regulatory β (CSK2B) subunits of protein kinase CK2 (Table 1 and Table S2), which is the primary kinase responsible for pre-phosphorylation of proteins prior to their pyrophosphorylation by 5-IP7 [11, 12, 14, 15].

**Table 1.** Selected interaction partners of IP6K1. The table indicates the total peptide numbers for each protein identified in replicate pull-downs of IP6K1 and GFP (negative control). Only known pyrophosphorylation substrate proteins, and the subunits of CK2 are shown.

| Protein ID /<br>(Gene ID) | Total peptide number |  |  |  |
| --- | --- | --- | --- | --- |
|  | SFB-IP6K1<br>Replicate 1 | SFB-IP6K1<br>Replicate 2 | SFB-GFP<br>Replicate 1 | SFB-GFP<br>Replicate 2 |
| AP3B1 / (AP3B1) | 9 | 7 | 0 | 0 |
| NOLC1 / (NOLC1) | 0 | 1 | 0 | 0 |
| TCOF / (TCOF1) | 1 | 17 | 0 | 0 |
| UBF1 (UBTF) | 3 | 0 | 0 | 0 |
| CSK21 / (CSNK2A1) | 9 | 8 | 0 | 0 |
| CSK22 / (CSNK2A2) | 3 | 0 | 0 | 0 |
| CSK2B / (CSNK2B) | 0 | 3 | 0 | 0 |

### 3.2. IP6K1 interacts with IP7 substrate proteins

To validate the binding of IP6K1 to 5-IP7 pyrophosphorylation substrate proteins, we expressed tagged IP6K1 in HEK293T cells, and performed co-immunoprecipitation analyses. V5-tagged IP6K1 was able to pull-down overexpressed AP3B1 (Figure 2A). Similarly, overexpressed NOLC1, a highly pyrophosphorylated nucleolar protein [11, 17], also co-precipitated with V5-tagged IP6K1 (Figure 2B). We confirmed the interaction of IP6K1 with another highly pyrophosphorylated 5-IP7 substrate, TCOF [17], which was detected in the IP6K1 interactome (Table 1). Endogenous TCOF was bound to SFB-tagged IP6K1 enriched on streptavidin-sepharose beads (Figure 2C). SFB-tagged IP6K1 was also able to bind endogenous UBF1, another nucleolar 5-IP7 pyrophosphorylation substrate protein [17]; Figure 2D). We have earlier shown that the oncoprotein MYC is a substrate for pyrophosphorylation by 5-IP7 synthesized by IP6K1 [14]. Although MYC was not present in the IP6K1 interactome in HEK293T cells, SFB-tagged IP6K1 was able to pull-down co-overexpressed MYC (Figure 2E). In summary, we observed the binding of IP6K1 to several 5-IP7 pyrophosphorylation substrate proteins – AP3B1, NOLC1, UBF1, TCOF and MYC.

**Figure 2:**
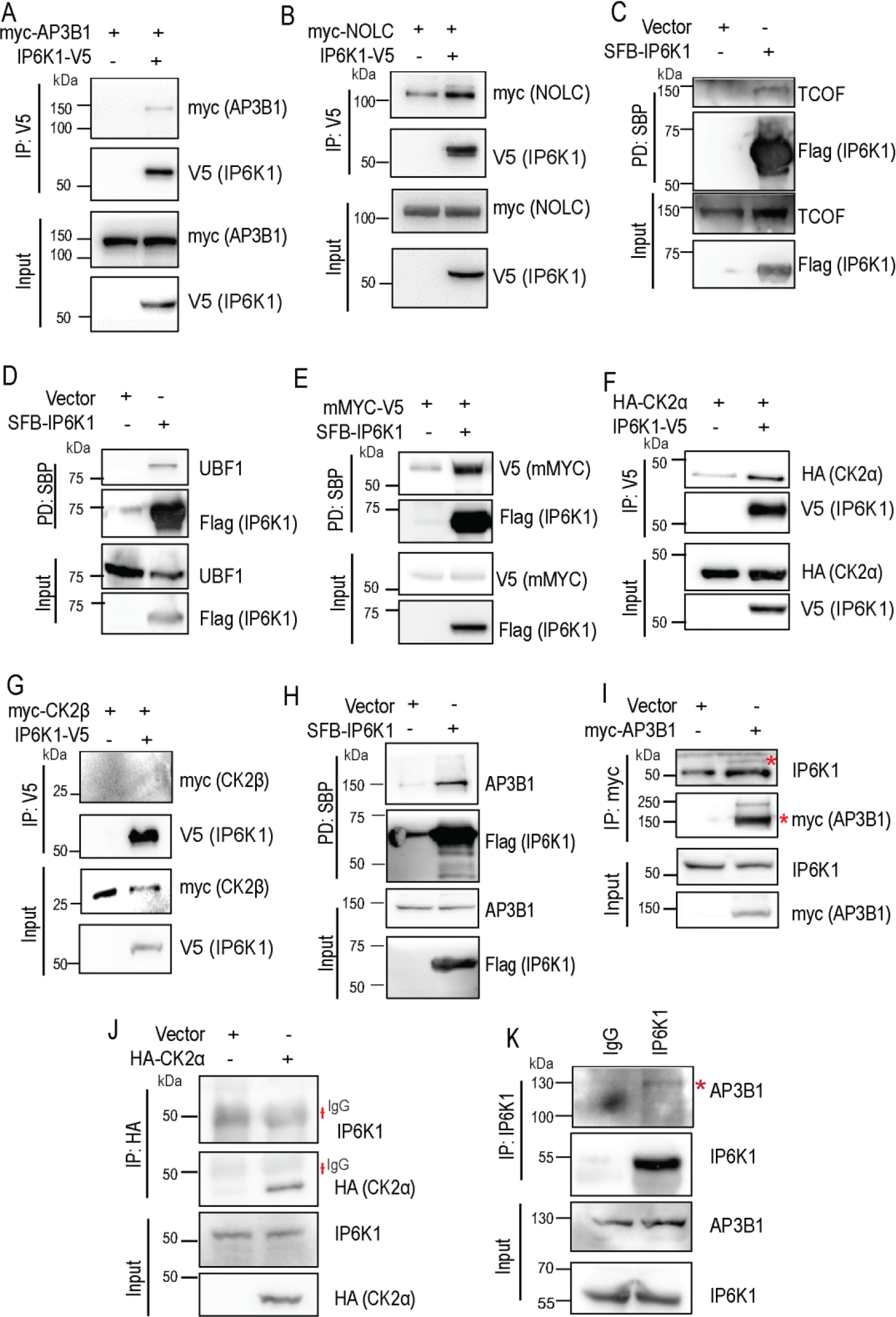
IP6K1 binds pyrophosphorylation substrate proteins and CK2. **(A-B)** Representative immunoblots examining co-precipitation of AP3B1 (A) and NOLC1 (B) with IP6K1. C-terminally V5-tagged IP6K1 and N-terminally myc-tagged AP3B1 or NOLC1 were transiently co-expressed in HEK293T cells, immunoprecipitated with an anti-V5 antibody, and probed to detect the myc epitope (N=3) for AP3B1; N=2 for NOLC1). **(C-D)** Representative immunoblots examining co-precipitation of endogenous TCOF (C) or UBF1 (D) with IP6K1. N-terminally SFB-tagged IP6K1 was transiently expressed in HEK293T cells, pulled down with streptavidin-sepharose beads, and probed to detect endogenous TCOF (N=3) or UBF1 (N=3). **(E)** Representative immunoblots examining co-precipitation of mouse MYC protein with IP6K1. N-terminally SFB-tagged IP6K1 and C-terminally V5-tagged MYC were transiently co-expressed in HEK293T cells, pulled down with streptavidin-sepharose beads and probed to detect the V5 tag (N=3). **(F)** Representative immunoblots examining co-immunoprecipitation of CK2α with IP6K1. C-terminally V5-tagged IP6K1 and N-terminally HA-tagged CK2α were transiently co-expressed in HEK293T cells, immunoprecipitated with an anti-V5 antibody, and probed to detect the HA tag (N=3). **(G)** Representative immunoblots examining co-immunoprecipitation of CK2β with IP6K1. C-terminally V5-tagged IP6K1 and N-terminally myc-tagged CK2β were transiently co-expressed in HEK293T cells, immunoprecipitated with an anti-V5 antibody, and probed to detect the myc epitope (N=2). CK2β could not be detected in the precipitate. **(H)** Representative immunoblots examining co-precipitation of endogenous AP3B1 with IP6K1. N-terminally SFB-tagged IP6K1 was transiently overexpressed in HEK293T cells, pulled down with streptavidin-sepharose beads, and probed to detect endogenous AP3B1 (N=3). **(I)** Representative immunoblots examining co-immunoprecipitation of endogenous IP6K1 with AP3B1. N-terminally myc-tagged AP3B1 was transiently expressed in HEK293T cells, immunoprecipitated with an anti-myc antibody, and probed to detect endogenous IP6K1 (N=2). **(J)** Representative immunoblots examining co-precipitation of endogenous IP6K1 with CK2α. N-terminally HA-tagged CK2α was transiently overexpressed in HEK293T cells, immunoprecipitated with an anti-HA antibody and probed to detect endogenous IP6K1 (N=3). IP6K1 could not be detected in the precipitate. **(K)** Representative immunoblots of the interaction between endogenous IP6K1 and endogenous AP3B1. HEK293T lysate was subjected to immunoprecipitation using an antibody against the N-terminal region of IP6K1, and probed for endogenous AP3B1 (N=3). Pre-immune IgG was used as a control. A red dagger symbol (†) corresponds to the IgG band and a red asterisk (*) marks the specific band of interest. An anti-Flag antibody was used to detect SFB-tagged proteins. Where indicated, cells were transfected with an empty vector as a control.

Next, we confirmed the interaction of IP6K1 with protein kinase CK2. Human CK2 is a heterotetramer consisting of two subunits, the catalytic α-subunit (sometimes replaced by the α’-subunit) and the regulatory β-subunit [30]. V5-tagged IP6K1was able to pull-down co-overexpressed HA-tagged CK2 α-subunit (CK2α) (Figure 2F). Although we had obtained a few peptides for the regulatory CK2 β-subunit in the IP6K1 interactome (Table 1), we were unable to detect the presence of overexpressed CK2 β-subunit in the IP6K1 pull-down (Figure 2G).

Lastly, we examined the interaction of endogenous proteins. Endogenous AP3B1 could be captured with SFB-tagged IP6K1 in HEK293T cells (Figure 2H). In the reverse direction, myc-tagged AP3B1 was able to interact with endogenous IP6K1 (Figure 2I). However, we were unable to detect any binding of endogenous IP6K1 to HA-tagged CK2α (Figure 2J), suggesting that the interaction between IP6K1 and CK2 may be transient. We confirmed the interaction between endogenous IP6K1 and AP3B1 - an antibody directed against the N-terminus of IP6K1 was able to co-precipitate endogenous AP3B1 in HEK293T cells (Figure 2K).

### 3.3. IP6K1 binds directly to AP3B1 and indirectly to CK2

The results described above suggest the formation of a protein complex by the 5-IP7 synthesizing enzyme IP6K1, the 5-IP7 pyrophosphorylation substrate protein AP3B1, and the pre-phosphorylating protein kinase CK2. It has earlier been shown that CK2 mediated pre-phosphorylation of AP3B1 can prime it for pyrophosphorylation by 5-IP7 synthesized by IP6K1 [12]. To check which of the interactions are direct and which are indirect, we co-overexpressed IP6K1, AP3B1, and CK2α. We examined the expression level and extent of co-precipitation of AP3B1 or CK2α with IP6K1 when these proteins were expressed independently or together. IP6K1 was able to pull-down both CK2α and AP3B1, independently, and when these proteins were co-expressed (Figure 3A). There was a marginal increase in the level of CK2α when it was co-overexpressed with AP3B1 (Figure 3A, B). However, when corrected for this increase in expression, the extent of binding of CK2α to IP6K1 was enhanced ∼3-fold when AP3B1 was co-expressed, suggesting that CK2α may bind to IP6K1 via AP3B1 (Figure 3A, B). Interestingly, the expression of AP3B1 increased by ∼2.6 fold when CK2α was co-expressed, and IP6K1 expression was held constant across both samples (Figure 3A, C). This suggests a role for protein kinase CK2 in regulating the cellular level of AP3B1. When normalized for increased AP3B1 levels, there was no further effect of the presence of CK2α on the binding of AP3B1 with IP6K1, suggesting a direct interaction between AP3B1 and IP6K1, independent of CK2 (Figure 3A, C). To verify this finding, we utilized GST-tagged recombinant IP6K1 expressed in *E. coli*. GST-IP6K1 immobilized on glutathione-sepharose beads was incubated with HEK293T cell lysates in which CK2α or AP3B1 were expressed independently or together. GST-IP6K1 was able to bind AP3B1 and CK2α independently, but the extent of binding with CK2α was significantly enhanced in the presence of AP3B1 (Figure 3D). No binding of either protein was seen with the GST control. As seen in the case of IP6K1 immunoprecipitated from HEK293T cells (Figure 3A, C), GST-IP6K1 purified from *E. coli* showed no difference in binding to AP3B1 in the presence or absence of CK2α, when normalized for the increased expression of AP3B1 in cells co-expressing CK2α (Figure 3D).

**Figure 3.**
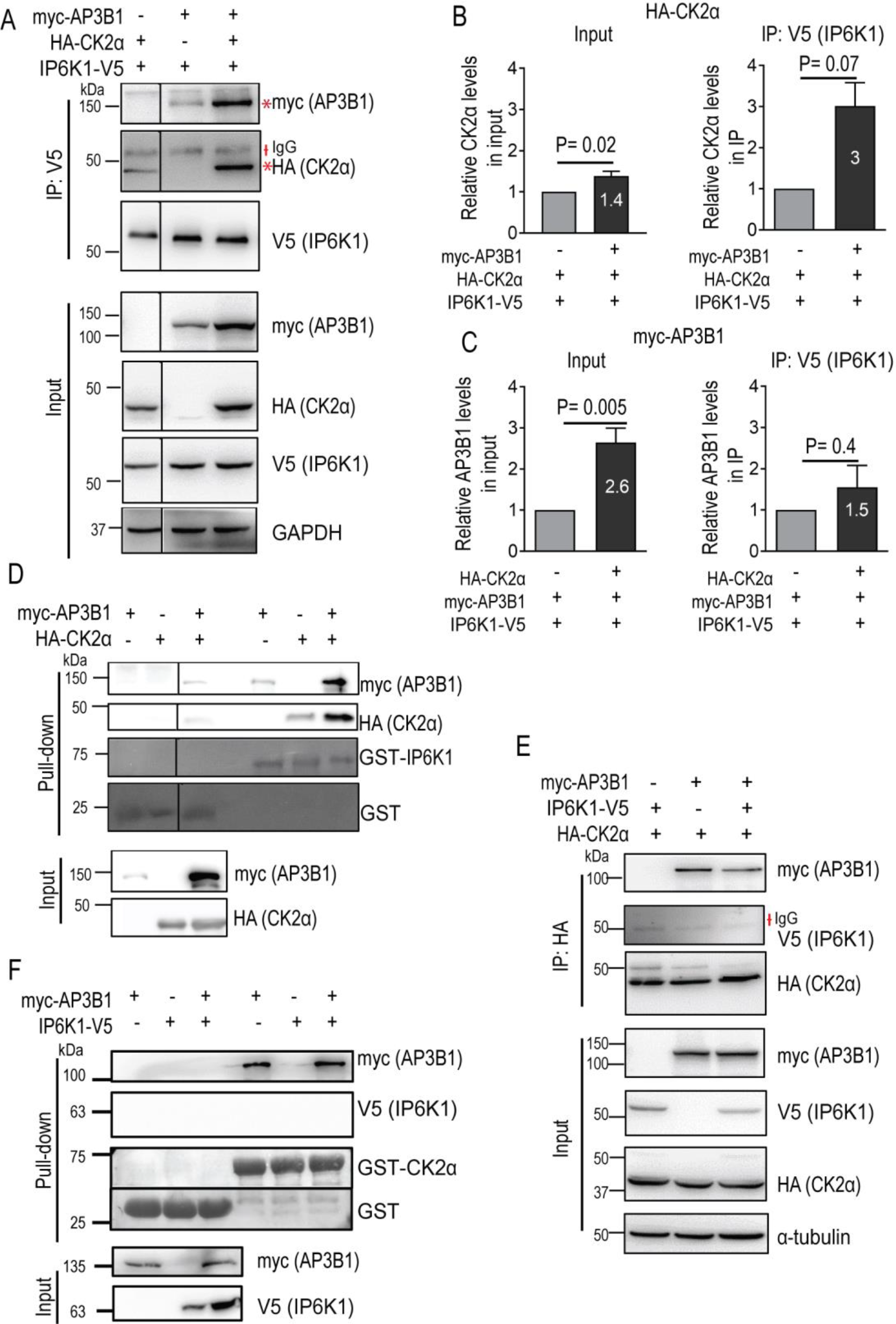
IP6K1 binds CK2 via AP3B1. **(A)** Representative immunoblots showing the effect of CK2α and AP3B1 co-expression on the extent of pull-down by IP6K1. V5-tagged IP6K1 was transiently co-expressed with HA-tagged CK2α and/or myc-tagged AP3B1 in HEK293T cells. Lysates were subjected to immunoprecipitation with an anti-V5 antibody and probed to detect the HA or myc epitopes. **(B)** Quantification of HA-CK2α expression level (Input) and extent of co-immunoprecipitation with IP6K1-V5 (IP) in cell lysates from (A) in the presence or absence of co-expressed myc-AP3B1. Data (mean ± SEM) were analyzed using a one-sample *t*-test (N=6 for Input and N=3 for IP). **(C)** Quantification of myc-AP3B1 expression level (Input) and extent of co-immunoprecipitation with IP6K1-V5 (IP) in cell lysates from (A) in the presence or absence of co-expressed HA-CK2α. Data (mean ± SEM) were analyzed using a one-sample *t*-test (N=6 for Input and N=3 for IP). **(D)** Representative immunoblots showing pull-down of AP3B1 and CK2α by purified IP6K1. GST or GST-IP6K1 purified from E. coli and immobilized on glutathione-sepharose beads was incubated with lysates from HEK293T cells transfected with HA-tagged CK2α and/or myc-tagged AP3B1. HA- and myc-tagged proteins were detected by immunoblotting and GST-tagged proteins were detected by Ponceau S staining (N=3). **(E)** Representative immunoblots showing the effect of AP3B1 and IP6K1 co-expression on the extent of pull-down by CK2α. HA-tagged CK2α was transiently co-expressed with myc-tagged AP3B1 and/or V5-tagged IP6K1 in HEK293T cells. Lysates were subjected to immunoprecipitation with an anti-HA antibody and probed to detect the V5 or myc epitopes (N=3). IP6K1 could not be detected in the precipitate. **(F)** Representative immunoblots examining pull-down of AP3B1 and IP6K1 by purified CK2α. GST or GST-CK2α purified from E. coli and immobilized on glutathione-sepharose beads was incubated with lysates from HEK293T cells transfected with V5-tagged IP6K1 and/or myc-tagged AP3B1. V5- and myc-tagged proteins were detected by immunoblotting and GST-tagged proteins were detected by Ponceau S staining (N=3). A red dagger symbol (†) corresponds to the IgG band and a red asterisk (*) marks the specific band of interest. The vertical lines in (A) and (D) indicate the removal of nonessential lanes from a single original blot for better depiction; the horizontal black line in (F) indicates cropping of the intervening region in the same blot.

As CK2α appears to bind IP6K1 via AP3B1 (Figure 3A, B, D), we wondered whether co-expression of AP3B1 would enhance the detection of IP6K1 in a pull-down of CK2α. However, as seen earlier in the absence of AP3B1 co-expression (Figure 2J), we could not detect any IP6K1 signal in the CK2α immunoprecipitate even in the presence of AP3B1 (Figure 3E). We then expressed CK2α fused with GST in *E. coli*, and incubated this protein with HEK293T cell extracts expressing IP6K1 or AP3B1 independently or together. Whereas GST-CK2α showed robust binding to AP3B1, we did not detect any IP6K1 in the pull-down (Figure 3F). Although CK2α expressed in HEK293T cells is seen to bind weakly with immobilized GST-IP6K1, the reverse binding of IP6K1 in HEK293T extracts with immobilized GST-CK2α, is not seen. One explantion for this observation is that the binding of CK2α to IP6K1 in the absence of co-expressed AP3B1 is indirect, and facilitated by other endogenous IP6K1 interacting partners in the cell extract. Together, these data suggest that in the tripartite protein complex formed by IP6K1, AP3B1, and CK2, IP6K1 binds directly with AP3B1, and indirectly with CK2 via AP3B1.

### 3.4. CK2 phosphorylates and regulates the level of AP3B1

Having observed an increase in the cellular level of AP3B1 when it is co-expressed with CK2α (Figure 3A, C), we next determined whether the protein kinase activity of CK2 is essential for this phenomenon. For this we utilized a catalytically inactive mutant of CK2α in which Lys68 is substituted with Met [31]. The level of AP3B1 increased upon co-expression of active CK2α (Figure 4A, C). However, co-expression of inactive CK2α had no effect on AP3B1 expression (Figure 4B, C), confirming that CK2α regulates AP3B1 via its kinase activity. To validate the phosphorylation of AP3B1 by CK2, we immunoprecipitated myc-tagged AP3B1 and incubated it with CK2 holoenzyme in the presence of radiolabeled [γ-^32^P]ATP. We observed robust phosphorylation of AP3B1 by CK2 (Figure 4D).

**Figure 4.**
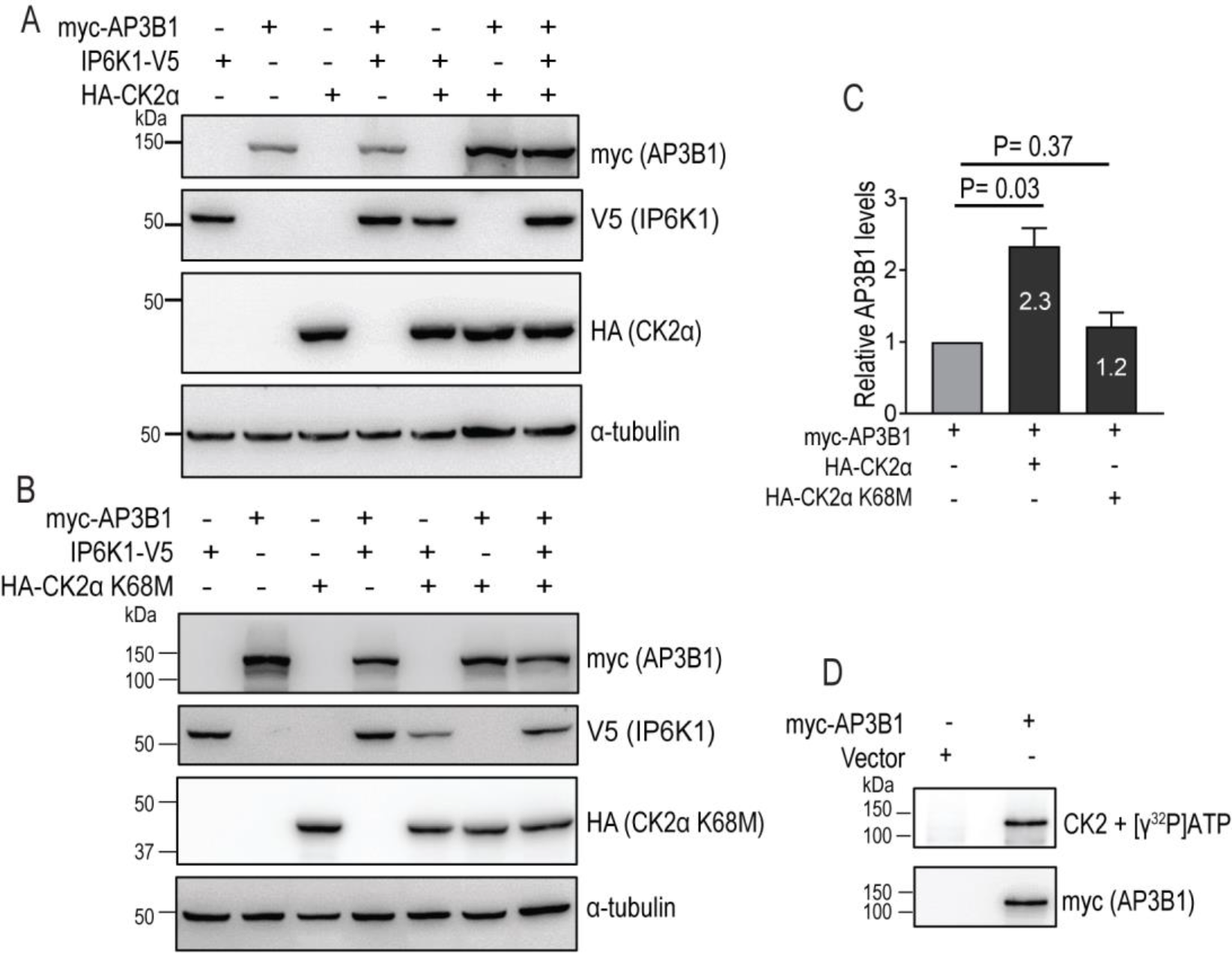
CK2 phosphorylates and elevates AP3B1 levels. **(A-B)** Representative immunoblots showing the effect of CK2 activity on the level of AP3B1. Myc-tagged AP3B1, V5-tagged IP6K1, and active (A) or catalytically inactive Lys68Met (B) HA-tagged CK2α were expressed in HEK293T cells individually or in combination. The immunoblots were probed to detect myc, V5 and HA epitopes. **(C)** Quantification of (A) and (B), where the bar graph represents mean fold change ± S.E.M. in the levels of myc-AP3B1, when transiently co-expressed with native CK2α or inactive CK2α K68M, in the absence of IP6K1. Data were analyzed using a one-sample *t*-test (N=3). **(D)** Phosphorylation of AP3B1 by CK2. Myc-AP3B1 immunoprecipitated from HEK293T cells was incubated with CK2 holoenzyme in the presence of radiolabeled [γ-^32^P]ATP. Images show autoradiography to determine phosphorylation (top) and immunoblotting with a myc-tag antibody (bottom). Cells transfected with an empty-vector were used as a negative control (N=3).

### 3.5. IP6K1 binds AP3B1 via an intrinsically disordered region in its N-lobe

The predicted structure of IP6K1 is broadly divided into amino (N-) terminal and carboxyl (C-) terminal lobes [32] (https://alphafold.ebi.ac.uk/entry/Q92551), and contains an intrinsically disordered region (IDR) within each lobe (IUPred3; https://iupred3.elte.hu/). Within the C-lobe lie the PDKG motif and IP helices, which are conserved across the family of IP kinases, and are involved in binding with the substrate IP6 [32]. We generated expression constructs corresponding to different fragments of IP6K1 spanning the N-lobe, C-lobe, a segment spanning the PDKG motif and IP helices, and intrinsically disordered regions designated IDR-1 and IDR-2, that lie within the N- and C-lobes respectively (Figure 5A). SFB-tagged IP6K1 fragments were co-overexpressed with AP3B1, and subjected to pull-down using streptavidin-sepharose beads. We observed maximum binding of AP3B1 with IP6K1 fragments corresponding to the N-lobe, IDR-1, and IDR-1+C-lobe (Figure 5B), suggesting that IP6K1 primarily interacts with AP3B1 via the long disordered region within its N-lobe. To confirm this, we monitored the effect of IDR-1 expression on the interaction between full length IP6K1 and AP3B1. The extent of co-precipitation of AP3B1 with IP6K1 was reduced by ∼50% in the presence of IDR-1 (Figure 5C, D), suggesting that IDR-1 expression has a dominant negative effect on the binding of IP6K1 with AP3B1.

**Figure 5:**
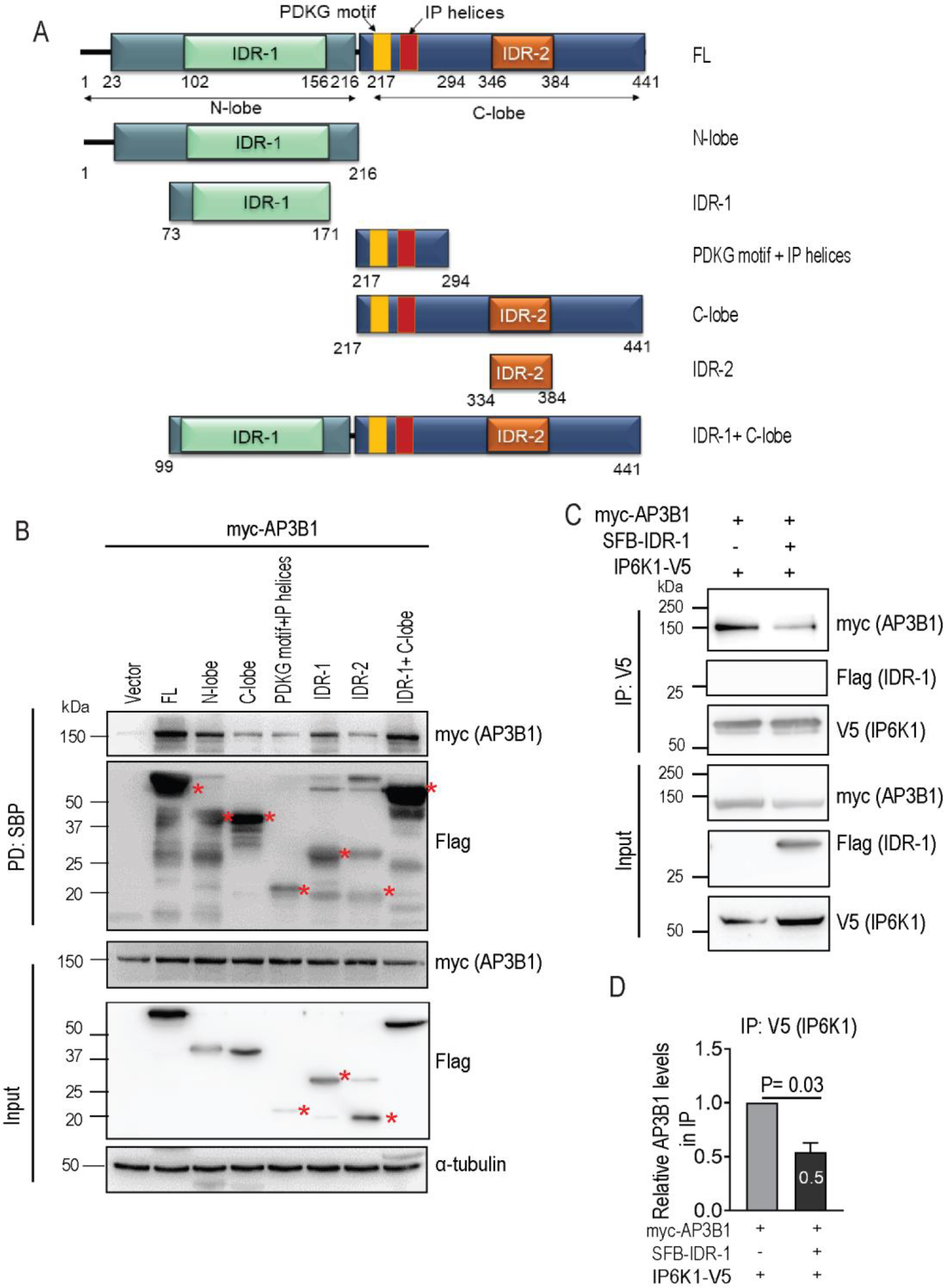
IP6K1 binds AP3B1 via its IDR-1 motif. **(A)** Domain map of human IP6K1 fragments used in this study. The first and last amino acid residue numbers of each construct are indicated. **(B)** Immunoblots examining the pull-down of AP3B1 with IP6K1 fragments. SFB-tagged IP6K1 fragments, co-expressed in HEK293T cells with myc-tagged AP3B1, were pulled down with streptavidin-sepharose beads and probed to detect the myc epitope. Pulled down IP6K1 fragments were probed using a Flag antibody (red asterisks (*) mark the specific bands of interest) (N=4). **(C)** Representative immunoblots showing the effect of IDR-1 overexpression on IP6K1-AP3B1 interaction. V5-tagged IP6K1 and myc-tagged AP3B1 were co-expressed in HEK293T cells in the absence or presence of SFB-tagged IDR-1 fragment of IP6K1, immunoprecipitated with an anti-V5 antibody, and probed to detect the myc epitope (N=3). **(D)** Quantification of the extent of co-immunoprecipitation of AP3B1 with IP6K1, in the absence or presence of SFB-IDR-1 in (C). Data (mean ± SEM) were analyzed using a one sample one-sample *t*-test (N=3).

### 3.6. Binding of IP6K1 to AP3B1 assists its pyrophosphorylation in vivo

Finally, we monitored how formation of a protein complex between the 5-IP7 synthesizing enzyme IP6K1 and the 5-IP7 pyrophosphorylation substrate AP3B1 affects intracellular pyrophosphorylation. AP3B1 has been shown to be pyrophosphorylated by radiolabeled 5-IP7 within its hinge region (residues 576-902) [12]. To establish that AP3B1 is indeed pyrophosphorylated by 5-IP7 synthesized by IP6K1 inside mammalian cells, we conducted a ‘back-pyrophosphorylation’ assay [26]. This technique compares the extent of in vitro pyrophosphorylation of a substrate protein isolated from cells with differential intracellular levels of 5-IP7. A target protein isolated from cells with lower 5-IP7 levels would exist in a hypopyrophosphorylated state compared with the same protein isolated from cells with higher 5-IP7 levels. Therefore, in the presence of radiolabeled 5-IP7, the hypopyrophosphorylated protein obtained from 5-IP7 deficient cells would accept a higher amount of radiolabeled phosphate compared with the protein isolated from 5-IP7 abundant cells. To conduct this assay, we relied on IP6K1 knockout HEK293T cells expressing either catalytically active or inactive (Lys226Ala) IP6K1. We observed an ∼80% decrease in 5-IP7 levels in cells expressing inactive IP6K1 compared with cells expressing the active form of the kinase (Figure 6A). AP3B1 over-expressed under these two conditions was immunoprecipitated and incubated with radiolabeled 5-IP7. There was a ∼2-fold higher extent of back-pyrophosphorylation on AP3B1 isolated from 5-IP7 deficient cells compared with AP3B1 from 5-IP7 sufficient cells (Figure 6B, C). This result firmly established that AP3B1 does indeed undergo intracellular pyrophosphorylation by 5-IP7 synthesized by IP6K1.

**Figure 6:**
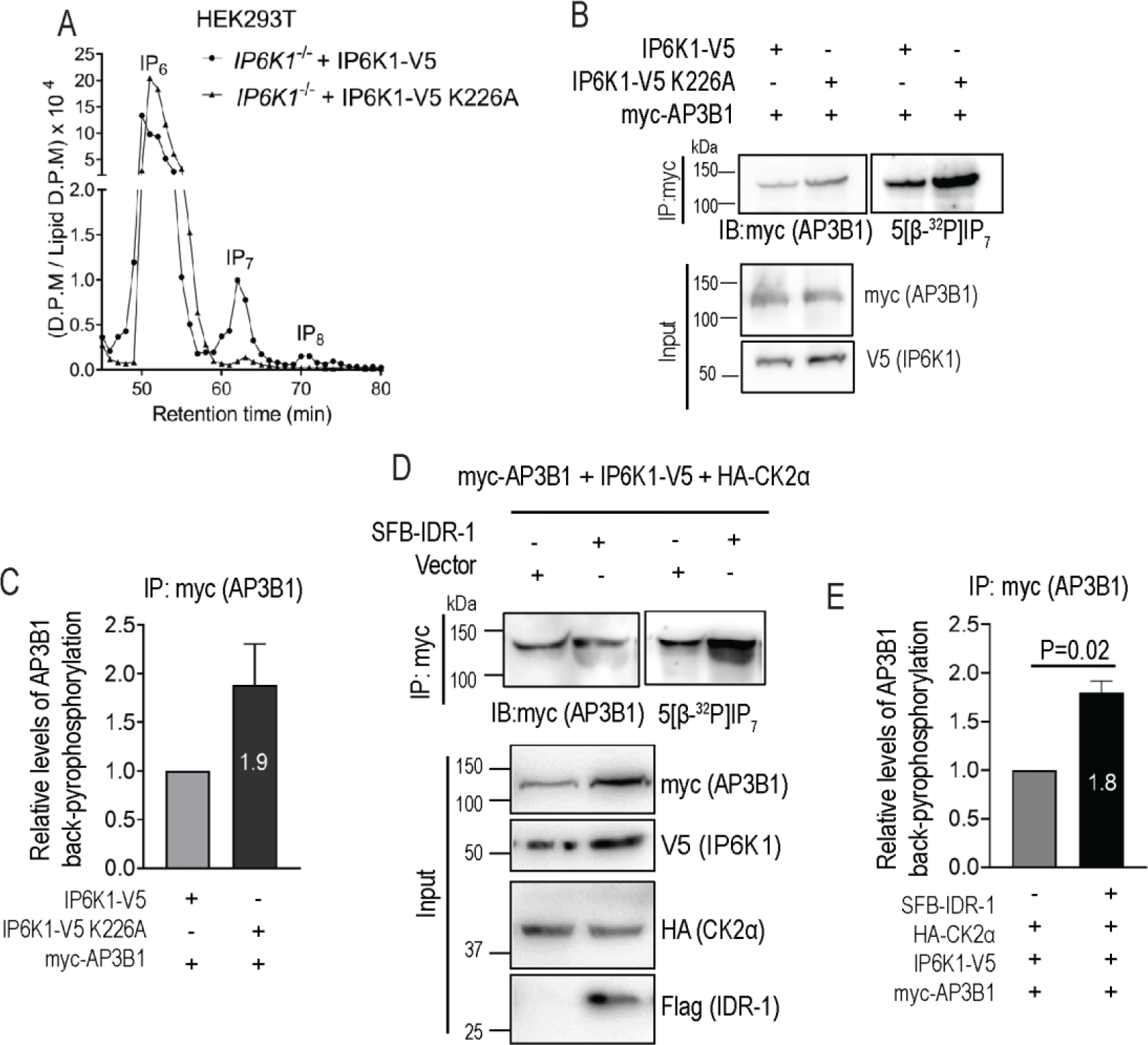
Pyrophosphorylation of AP3B1 is reduced by destabilizing protein complex formation. **(A)** HPLC profiles of [3H] inositol labelled IP6K1-/- HEK293T cells transiently overexpressing either active or catalytically inactive (Lys226Ala) V5-tagged IP6K1. Soluble inositol phosphate counts were normalized to the total lipid inositol count for each sample. Peaks corresponding to IP6, IP7, and IP8 are indicated. Data are representative of two independent experiments. **(B)** Back-pyrophosphorylation assay to demonstrate in vivo AP3B1 pyrophosphorylation. Myc-tagged AP3B1 co-expressed with either active or inactive V5-tagged IP6K1 in IP6K1-/- HEK293T cells was immunoprecipitated, incubated with 5[β-^32^P]IP7, resolved on a 4-12% NuPAGE Bis-Tris gel and transferred to a PVDF membrane. Representative images show immunoblotting to detect the myc epitope (left) and autoradiography to detect pyrophosphorylation (right). Input levels of all the co-expressed proteins are shown (N=2). **(C)** Quantification of (B) showing the fold change (mean ± range) in the extent of back-pyrophosphorylation of myc-AP3B1, normalized to its immunoprecipitated protein levels, when co-expressed with inactive vs active IP6K1 (N=2). **(D)** Back-pyrophosphorylation assay to monitor effect of IDR-1 on in vivo AP3B1 pyrophosphorylation. myc-tagged AP3B1, co-expressed in IP6K1-/- HEK293T cells with V5-tagged IP6K1 and HA-tagged CK2α, in the presence or absence of SFB-tagged IDR-1, was immunoprecipitated with myc-tag antibody and subjected to pyrophosphorylation in presence of 5[β-^32^P]IP7 as in (B). Representative images show immunoblotting to detect the myc epitope (left) and autoradiography to detect pyrophosphorylation (right). Input levels of all the co-expressed proteins are shown (N=3). **(E)** Quantification of (D) showing the fold change in the extent of back-pyrophosphorylation of myc-AP3B1, normalized to its immunoprecipitated protein levels, when expressed in the absence or presence of IDR-1. Data (mean ± SEM) were analyzed using a one-sample *t*-test (N=3).

Next, we relied on a similar back-pyrophosphorylation method to determine whether the binding between IP6K1 and AP3B1 aids intracellular pyrophosphorylation of AP3B1. For this, we co-overexpressed IP6K1, AP3B1 and CK2α in the presence or absence of the IP6K1 IDR1 fragment. Co-expression of IDR1 would decrease the intracellular interaction between AP3B1 and full length IP6K1 (Figure 5C, D), and would therefore reduce the local synthesis of 5-IP7 in the vicinity of AP3B1. We observed a ∼2-fold increase in back-pyrophosphorylation of AP3B1 isolated from cells that over-expressed SFB-IDR-1 along with IP6K1, compared with AP3B1 from control cells that expressed IP6K1 alone (Figure 6D, E). This data confirms that the dominant negative effect of IDR-1 on formation on the IP6K1-AP3B1 complex hinders in vivo pyrophosphorylation of AP3B1.

## 4. Discussion

Pyrophosphorylation of proteins by donating their β-phosphate moiety to a pre-phosphorylated serine residue is a mechanism by which PP-IPs modulate protein function. Although this unique enzyme-independent post-translational modification has been studied for some years, intracellular processes that influence protein pyrophosphorylation are not known. Here, we provide proof that the 5-IP7 synthesizing enzyme IP6K1 binds to several proteins that are known to undergo 5-IP7 mediated pyrophosphorylation. In the case of the pyrophosphorylation substrate AP3B1, we demonstrate its direct binding to a disordered region in the N-lobe of IP6K1, and to the α-subunit of protein kinase CK2. Phosphorylation of serine residues on AP3B1 by CK2 would prime them for pyrophosphorylation. The local synthesis of 5-IP7 by IP6K1 bound to AP3B1 would increase the relative abundance of 5-IP7 over IP6, and promote mass action driven pyrophosphorylation of the prephosphorylated serine residues on AP3B1 (Figure 7). In this way, formation and dissolution of a tripartite complex between the pyrophosphorylation substrate, the priming protein kinase, and the 5-IP7 synthesizing IP6 kinase, would provide cells with a mechanism to control enzyme-independent serine pyrophosphorylation by 5-IP7.

**Figure 7:**
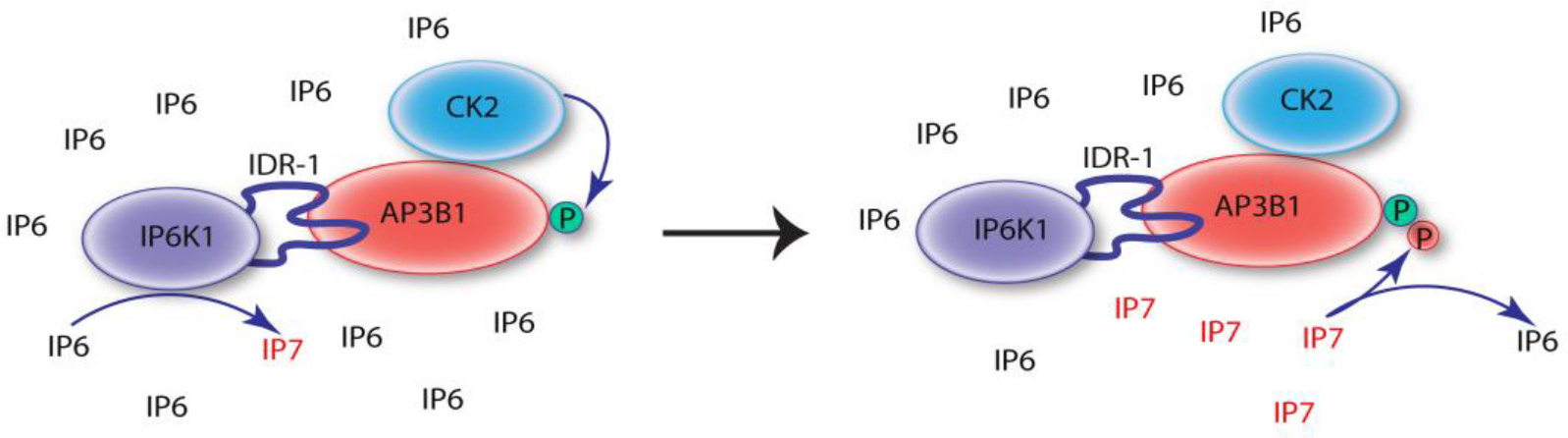
Formation of a protein complex facilitates 5-IP7 mediated pyrophosphorylation. The α subunit of protein kinase CK2 binds to AP3B1 and catalyzes its phosphorylation, priming it for 5-IP7-mediated pyrophosphorylation. IP6K1 interacts with AP3B1 via its IDR-1 region and synthesizes 5-IP7 in the vicinity of the complex. A local increase in the concentration of 5-IP7 promotes the mass-action driven transfer of its β-phosphate moiety to pyrophosphorylate AP3B1.

Pyrophosphorylation occurs predominantly on Ser residues that lie in the vicinity of acidic Asp and Glu residues, within an intrinsically disordered region of the substrate protein [9, 17]. Given the sequence context of the target Ser, its priming phosphorylation is usually catalyzed by acidophilic protein kinases such as CK2 [9, 14, 15, 17]. Although an enzyme is not needed to catalyze the phosphotransfer from a PP-IP to a prephosphorylated Ser (phospho-Ser), the presence of excess Mg^2+^ is essential for the reaction [9]. It is likely that Mg^2+^ ions coordinate the PP-IP with Asp and Glu residues near the target phospho-Ser to bridge the two ‘substrates’ in the absence of a classical ‘enzyme’. Intrinsic disorder in the surrounding region may play an important role in facilitating access of the target phospho-Ser to the Mg^2+^-PP-IP complex [9]. The data presented here show that a disordered region in the N-lobe of IP6K1 binds AP3B1. This region of intrinsic disorder is present in all three IP6K isoforms, and is the most variable sequence between them, possibly conferring differential binding specificity to the individual IP6Ks. Disordered sequences are known to provide scaffolds that facilitate protein-protein interaction [33, 34]. Binding of the IDR regions in IP6Ks with the disordered pyrophosphorylation substrate sequence could be a conserved mechanism to bring these proteins together and facilitate 5-IP7-mediated pyrophosphorylation.

One additional problem presented by the absence of an enzyme to catalyze pyrophosphorylation is that the target sequence would just as easily complex with Mg^2+^-IP6 as it would with Mg^2+^-PP-IP. Indeed, pyrophosphorylation by 5-IP7 is inhibited in the presence of excess IP6 [10]. A majority of proteins that bind IP6 are also shown to bind IP7 [8, 19, 35, 36]. In most organisms and cell types studied till date, the intracellular abundance of IP6 exceeds that of PP-IPs, although there are some notable exceptions, including the slime mould *Dictyostelium discoideum* in which IP7 and IP8 levels are similar to those of IP6 [37, 38]. A recent analysis of inositol phosphates in mouse tissues and human cell lines using the highly sensitive capillary electrophoresis electrospray ionization mass spectrometry method has confirmed that mammals have a 5-50 fold excess of IP6 over 5-IP7 [20]. In this light, given that transfer of the β-phosphate from a PP-IP to a phospho-Ser is likely a reaction driven by crowding of reactants, the relative local abundance of the donor PP-IP over IP6 (or even IP5) would be a factor controlling the extent of intracellular protein pyrophosphorylation. Our data showing that IP6K1 binds to many known pyrophosphorylated proteins suggests that local synthesis of 5-IP7 by an IP6 kinase bound to the pyrophosphorylation substrate protein may be a general mechanism to overcome the problem of excess IP6 over PP-IPs.

Although this work has extensively characterized the interaction between AP3B1, IP6K1 and CK2, a similar complex may be formed between other pyrophosphorylation substrates, IP6K1 (or other IP6 kinases) and CK2 (or other acidophilic protein kinases). We have shown that IP6K1 binds the nucleolar proteins NOLC1, TCOF and UBF1, and the nuclear oncoprotein MYC, all of which are documented substrates for pyrophosphorylation [14, 17]. IP6K1 has been shown to co-localize with UBF1 in the nucleolus, where many other pyrophosphorylated proteins are found [17]. In addition to these validated interactors, we note the presence of several other pyrophosphorylated proteins in the IP6K1 interactome (Table S2). CK2 is known to be expressed in most human cells and tissues (https://www.proteinatlas.org/ENSG00000113712-CSNK1A1), and is known to phosphorylate many proteins, primarily targeting Ser/Thr residues with an acidic Glu/Asp at the +3 position [39, 40]. Our data shows that CK2 phosphorylation increases the level of AP3B1. We do not yet know whether the phosphosites on AP3B1 responsible for this increase overlap with or are distinct from the sites that undergo subsequent pyrophosphorylation. Nevertheless, regulation of CK2 activity and subcellular localization may provide another handle for cells to control pyrophosphorylation. Interestingly, NOLC1 (also called Nopp140) has been shown to bind to CK2α and inhibit its catalytic activity, and IP6 binding to CK2α at the same site reverses this inhibition [41]. 5-IP7 synthesized by IP6K2 binds CK2 and promotes its kinase activity [27]. The regulation of CK2 activity by the binding of IP6 or 5-IP7, and competition with negatively charged protein substrates, would add another layer of complexity to the modulation of protein pyrophosphorylation by formation of a substrate-CK2-IP6K complex.

## Supplementary Materials

Table S1: Mass Spectrometry data, Table S2: CRAPome analysis, Table S3: DAVID Functional annotation clustering

## Author Contributions

Conceptualization, A.H., J.S.L., A.Sh., S.G, and R.B.; methodology, A.H., A.Sh., J.S.L., and S.G; validation, A.H., A.Sh., J.S.L., M.P., S.G., A.Si., and R.B.; formal analysis, A.H., J.S.L., and R.B.; investigation, A.H., A.Sh., J.S.L., M.P., S.G., and A.Si.; writing—original draft preparation, A.H., J.S.L., S.G., and R.B.; writing—review and editing, J.S.L., S.G., and R.B.; visualization A.H., J.S.L.; supervision, R.B; project administration, R.B.; funding acquisition, R.B. All authors have read and agreed to the published version of the manuscript.

## Data Availability Statement

The mass spectrometry data used in this manuscript have been deposited to the MassIVE, a full member of the Proteome Xchange Consortium and can be accessed through Data set identifier ID-MassIVE MSV000092218 (https://massive.ucsd.edu).

## Acknowledgments

We thank Dr. Solomon Snyder, Johns Hopkins School of Medicine, Baltimore, USA for human AP3B1 cDNA, Human CK2α cDNA, pGEX6P2-CK2α and pGEX6P1-IP6K1 plasmid; T. Mohanty and R. Manorama for generating radiolabeled IP7; Jayraj Sen for the generation of an antibody directed against the N terminus of IP6K1; Sitalakshmi Thampatty for the construction of SFB-GFP and SFB-IP6K1 expression plasmids; Vineesha Oddi for the construction of IP6K1-V5 expression plasmid. We also thank all members of the Laboratory of Cell Signalling for valuable feedback.

